# How many (distinguishable) classes can we identify in Single Particle Analysis?

**DOI:** 10.1101/2025.07.14.664719

**Authors:** O. Lauzirika, M. Pernica, D. Herreros, E. Ramírez-Aportela, J. Krieger, M. Gragera, M. Iceta, P. Conesa, Y. Fonseca, J. Jiménez, J. Filipovic, J.M. Carazo, C.O.S. Sorzano

## Abstract

Heterogeneity in cryoEM is essential for capturing macromolecule structural variability, reflecting their functional states and biological significance. However, estimating heterogeneity remains challenging due to particle misclassification and algorithmic biases, which can lead to reconstructions that blend distinct conformations or fail to resolve subtle differences. Furthermore, the low signal-to-noise ratio (SNR) inherent in cryo-EM data makes it nearly impossible to detect minute structural changes, as noise often obscures subtle variations in macromolecular projections. In this paper, we investigate the use of p-values associated with the null hypothesis that the observed classification differs from a random partition of the input dataset, thereby providing a statistical framework for determining the number of distinguishable classes present in a given dataset.

## 1 Introduction

In cryoEM, classification approaches can be broadly categorized as discrete or continuous (Sorzano et al., 2019; Tang et al., 2023; Toader et al., 2023; Kimanius and Schwab, 2024). Discrete classification aims to group particles into distinct, non-overlapping classes, each representing a specific structural conformation of the macromolecule. This method is ideal for resolving well-separated states, such as open and closed conformations, but struggles with more nuanced structural variability or when the number of distinct states is unclear. On the other hand, continuous classification models structural variability as a smooth continuum of conformations, better capturing gradual transitions or flexible regions within the macromolecule. However, continuous methods face challenges such as high computational demands and difficulties interpreting the results, particularly under the low signal-to-noise ratio (SNR) typical of cryoEM data. This paper focuses on discrete classification, designed to address cases of true discrete heterogeneity and approximate continuous heterogeneity by quantization. As highlighted in Sorzano et al. (2022), discrete classification can be highly unstable, especially in noisy datasets. Factors such as class imbalance, the “3D attraction effect,” and inconsistencies across repeated classifications often compromise the accurate identification of structural classes. These challenges underscore the importance of developing robust methodologies to enhance classification reliability and call for cautious interpretation of results. In this context, this paper examines the limitations of current approaches and explores potential strategies for improvement.

In Sorzano et al. (2022), we proposed that a practical way to identify parameter misestimates is to estimate the same parameter multiple times, preferably using different algorithms or, at the very least, through various executions of the same algorithm with non-deterministic variations, such as random initialization. For discrete 3D classification, this approach entails performing the classification repeatedly at each level. While this practice is rarely adopted due to its high computational cost, the demonstrated instability of 3D classification underscores the risk of basing biological interpretations solely on the results of a single execution, potentially leading to compromised conclusions.

Several works have addressed the problem of reliably estimating the number of classes and/or their composition:

- Rabuck-Gibbons et al. (2022) tackled the instability of 3D classification in cryo-EM datasets by introducing a workflow that iteratively divides an input dataset into two smaller subsets until the volume difference between the subclasses falls below a predefined threshold. Subsequently, similar maps are merged into larger classes through hierarchical clustering using their correlation as a similarity measure. However, this approach presents two limitations: determining an appropriate signal threshold for stopping the subdivision process is challenging, and the user must select the final number of clusters based on the dendrogram, which can be difficult to interpret, especially for complex datasets.
- In a similar vein, Gomez-Blanco et al. (2022) proposed a divisive hierarchical approach where, at each step, the dataset is split into 2 or 3 classes. The stopping criteria for further subdivision were: (1) the number of particles in a class must exceed a user-defined threshold (typically 5,000–10,000 particles), and (2) the resolution of the class, as determined by the classification method, must surpass the resolution predicted by the ResLog plot for the given particle count. If both conditions were satisfied, the class was further divided. Finally, to group similar maps, each map was represented by a feature vector derived from its truncated PCA representation, and clustering was performed using the Affinity Propagation algorithm, which measured distances in the feature vector space using the Euclidean metric.
- Zhou et al. (2022) tackled the problem by incrementally classifying the input dataset into *K* = 2, 3, 4, … classes. They measured the similarity between raw particle images and their corresponding reconstructed 3D references for each classification. The optimal number of classes was determined as the *K* that maximized the average variance of this similarity score within each class. The rationale is that an incorrect number of classes—either too few or too many—reduces the within-class variance. With too few classes, particles from different structural states are mixed, making it impossible for the representative model to accurately reflect any subset of particles. With too many classes, noise overfitting occurs, artificially increasing particle similarity within each class. As highlighted by Tibshirani et al. (2001), selecting the number of classes solely by inspecting within-cluster dispersion—such as minimizing intra-class variability—can be misleading and prone to substantial error, as it lacks a principled statistical basis. This concern is well recognized in the pattern recognition community, prompting the development of more statistically grounded approaches, such as the gap statistic.

A significant limitation of all these approaches is their reliance on a single classification step for each *K*, which inherently makes the results susceptible to the stochastic nature and instability of the classification process, particularly as *K* (the number of classes) increases. In Single Particle Analysis, classification instability arises due to noise, particle misalignments, and biases introduced by initial conditions or the specific algorithm. As *K* grows, separating particles into increasingly finer classes amplifies the risk of overfitting noise or misassigning particles to incorrect classes. This can result in artificial or noisy classes, blending of distinct conformations, or failure to identify low-populated conformational states.

In this article, we propose a divisive approach similar to the methods introduced by Rabuck-Gibbons et al. (2022) and Gomez-Blanco et al. (2022). Our approach uses binary subdivisions, as dividing into more classes can increase classification instability. However, unlike previous methods, we repeat the same subdivision multiple times to estimate their consistency. The stopping criterion is determined by the p-value associated with the null hypothesis that the observed subdivision is random rather than reflecting the genuine structural features of the images. This statistical foundation ensures that the divisions are driven by meaningful data characteristics rather than noise or arbitrary thresholds on the signal difference, the number of particles, or the class resolution. We are aware that performing multiple executions of the same classification step may be computationally tedious; however, at present, it remains the only practical way to estimate the uncertainty of the classification outcome. In the future, developers may introduce faster classification algorithms or alternative techniques to more efficiently quantify classification uncertainty.

## 2 Methods

Suppose we partition a set of *N* images into *K* classes and that the images truly exhibit a discrete class structure. In that case, we expect them to be consistently classified into stable groups, regardless of the number of classes used for the split, as illustrated in Fig. 1. If the classification algorithm successfully identifies the underlying classes, specific subsets of images should consistently remain grouped across different levels of the hierarchy. For instance, in the provided example, we would expect the images in Class 1 of Classification C to also be grouped in Class B1 and Class A1 at higher levels of the hierarchy. Conversely, if the classification algorithm fails to consistently separate the images, we would expect the images to be randomly distributed among the classes in subsequent classifications. This will be our null hypothesis. In this scenario, let 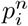 represent the proportion of images in the *i*-th class during the *n*-th classification. The expected proportion of images classified into Class *i*_1_ in the first classification, *i*_2_ in the second, and so on, up to *i*_*n*_ in the *n*-th classification, would be given by:

**Figure 1:**
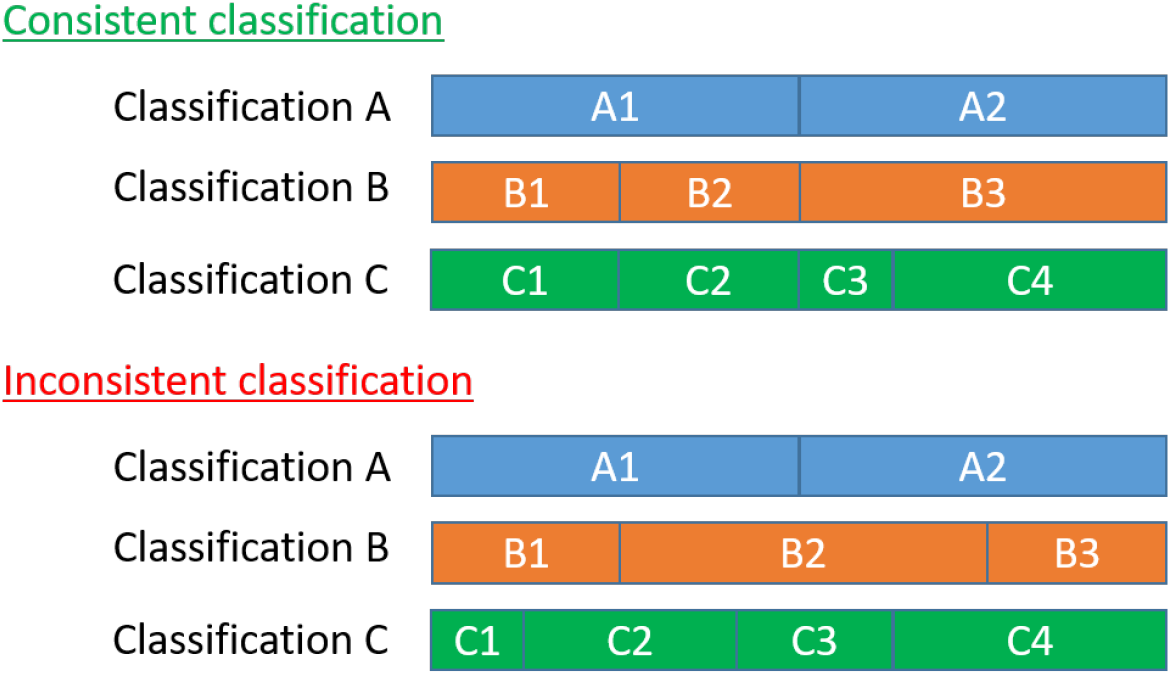
Examples of consistent and inconsistent classifications of a set of images into 2, 3, and 4 classes, named A, B, and C. In consistent classification, the grouping structure of the larger groups is respected, whereas in inconsistent classification, it is not.

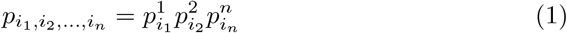

The expected size of this set can be modeled using a binomial distribution with parameters *N* and 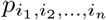. However, in our classification, which we aim to ensure is not random, we observe an actual size 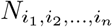. To assess whether this observed size is consistent with the null hypothesis of random classification, we calculate the probability of observing a size equal to or greater than 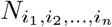 under the null hypothesis. This probability provides a p-value associated with each observed size. We reject the null hypothesis of random classification if this size is smaller than a given threshold (typically, 0.05, although we may lower it to 0.01 or even smaller).

It is essential to note that the reproducibility of a classification, as assessed by our method, does not guarantee its accuracy in itself. A classification can appear stable across repetitions not because it reflects the true structural heterogeneity of the dataset, but due to a persistent bias introduced by the underlying algorithm, its initialization, regularization strategies, or the weighting of specific image features. To address this limitation, the most robust application of our framework involves comparing classifications obtained from different algorithms. By incorporating cross-method comparisons into the core of the procedure, we aim to reduce the risk of algorithm-specific biases being mistaken for true structural features. While our statistical framework quantifies the internal consistency of the classification process, it does not detect systematic biases that may be shared across repeated runs of the same method. In this sense, the technique evaluates classification variability, not correctness. Complementary validation steps—such as inspecting the resulting 3D reconstructions, performing cross-method comparisons, or interpreting the results in light of prior biological knowledge—remain critical for assessing the plausibility and validity of the identified classes.

In our divisive algorithm, each input dataset is split into two classes. This process is repeated *n* times to identify subsets of images that are consistently and significantly grouped, as described above. These identified subsets are treated as new classes, which can then be further divided into smaller subsets. Images that cannot be consistently separated are combined into a new class and reintroduced into the algorithm. The algorithm continues branching until no further meaningful divisions can be made. At this point, that branch is terminated. While our choice of binary subdivision is arbitrary, the method is general and would apply equally to splits into three or more classes. However, as the number of output classes increases, the classification typically becomes less stable, making it harder to identify statistically significant and reproducible groupings. Therefore, binary splits provide a practical balance between expressive power and robustness.

Our divisive algorithm is flexible and independent of the specific classification algorithm used to separate the images. Several options can be employed: (1) a classification algorithm that uses previously estimated angular assignments, keeping them fixed while estimating only the class memberships; (2) a joint estimation of angular assignments and class memberships, which can be performed either as a local refinement (using prior estimates) or as a global refinement (without relying on prior estimates).

## 3 Results

We validated our algorithm on several representative datasets, utilizing a classification workflow that incorporated multiple steps as necessary. These steps might include map reconstruction using Fourier gridding techniques (Abrishami et al., 2015), local refinement of angular and translational parameters with Xmipp highres (Sorzano et al., 2018), and unsupervised 3D classification in CryoSparc without alignment.

### 3.1 Apoferritin

Apoferritin is a standard benchmark in cryo-electron microscopy (cryoEM) due to its structural stability and is frequently used for calibration. Given its homogeneity, we did not expect our method to identify multiple classes. We analyzed 17,000 polished particles from EMPIAR-11796 (Aiyer et al., 2024), using their Relion-derived pose assignments. The particles were classified into two groups across three independent CryoSparc runs without alignment. As anticipated, only one class consistently showed a statistically significant p-value, comprising 100% of the particles.

### 3.2 Ribosembly

The Ribosembly dataset, part of the CryoBench benchmark suite (Jeon et al., 2024), contains 335,240 synthetic cryo-EM particle images corresponding to 16 well-defined bacterial ribosome assembly intermediates. These intermediates share a conserved structural core and progressively incorporate additional proteins and rRNA, providing a realistic model of compositional heterogeneity during ribosome maturation. The dataset includes ground-truth atomic structures, density maps, and imaging parameters, making it well-suited for assessing the performance of classification algorithms.

We applied our divisive classification workflow iteratively until no further statistically significant subdivisions could be identified. Given the known angular assignments, each step involved reconstructing candidate subgroups using Relion’s Fourier gridding and performing three independent two-class classifications with CryoSparc, without alignment. We then derived consensus classes, computed their associated p-values, retained statistically significant splits, regrouped non-significant subsets, and continued the process on the remaining data.

Table 1 shows the distribution of particle images across the 16 ground truth classes in the Ribosembly dataset and their assignments after two iterations of our classification procedure. Although the workflow involved additional iterations, we limit the table to the first two levels for clarity. The results show that some ground truth classes are relatively well-preserved across classifications — for example, most particles from classes 3 and 14 consistently remain grouped. In contrast, other classes, such as 0 and 7, appear more dispersed. While this is not a strict rule, it is generally observed that large ground truth classes tend to remain cohesive. In contrast, smaller classes are more prone to fragmentation across output classes. It is essential to note that our algorithm does not introduce this dispersion; rather, it reflects the behavior of the underlying 3D classification method. Our algorithm groups particles that were consistently assigned together by the 3D classifier.

**Table 1:** Distribution of Ribosembly ground truth classes (shown as a percentage of the total number of images) and their assignment to selected classification outputs after multiple rounds of classification. Classes C1 and C2 follow the first statistically reliable classification. The numbers represent the percentage of the underlying true class images in each of the calculated classes. Classes C11, C12, C13, and C14 correspond to the statistically reliable split of class C1 into statistically distinguishable subsets. Similarly, classes C21 and C22 correspond to the statistically reliable split of class C2. The last row shows the purity of each class, which is computed as the maximum value in the column divided by the sum of all the values.

| True class | Total % | C1 | C2 | C11 | C12 | C13 | C14 | C21 | C22 |
| --- | --- | --- | --- | --- | --- | --- | --- | --- | --- |
| 0 | 2.7 | 87.1 | 12.7 | 15.3 | 68.0 | 2.4 | 1.4 | 10.0 | 2.6 |
| 1 | 4.3 | 93.8 | 6.1 | 5.7 | 87.3 | 0.5 | 0.3 | 4.4 | 1.7 |
| 2 | 7.0 | 95.4 | 4.5 | 2.4 | 92.4 | 0.4 | 0.2 | 3.3 | 1.2 |
| 3 | 13.2 | 99.2 | 0.8 | 98.9 | 0.1 | 0.1 | 0.1 | 0.6 | 0.2 |
| 4 | 9.1 | 99.5 | 0.5 | 99.4 | 0.0 | 0.0 | 0.1 | 0.4 | 0.1 |
| 5 | 11.5 | 98.8 | 1.1 | 0.7 | 0.2 | 97.6 | 0.4 | 0.9 | 0.3 |
| 6 | 1.2 | 99.7 | 0.3 | 15.0 | 0.0 | 0.1 | 83.7 | 0.3 | 0.0 |
| 7 | 1.2 | 22.8 | 76.9 | 22.0 | 0.1 | 0.3 | 0.4 | 75.8 | 1.1 |
| 8 | 3.8 | 0.5 | 99.5 | 0.3 | 0.0 | 0.0 | 0.2 | 99.5 | 0.1 |
| 9 | 3.6 | 99.9 | 0.1 | 0.3 | 0.0 | 0.1 | 99.5 | 0.1 | 0.0 |
| 10 | 5.4 | 99.4 | 0.6 | 0.2 | 0.0 | 0.1 | 99.0 | 0.6 | 0.0 |
| 11 | 1.5 | 88.7 | 11.1 | 0.6 | 0.0 | 0.2 | 87.8 | 6.3 | 4.7 |
| 12 | 10.5 | 0.1 | 99.9 | 0.0 | 0.0 | 0.0 | 0.0 | 99.9 | 0.0 |
| 13 | 11.2 | 0.1 | 99.9 | 0.0 | 0.0 | 0.0 | 0.1 | 99.9 | 0.0 |
| 14 | 12.1 | 0.4 | 99.6 | 0.1 | 0.0 | 0.0 | 0.2 | 0.1 | 99.5 |
| 15 | 1.7 | 1.5 | 98.4 | 0.6 | 0.0 | 0.2 | 0.7 | 0.3 | 98.1 |
| Purity |  | 0.10 | 0.16 | 0.38 | 0.37 | 0.96 | 0.27 | 0.25 | 0.48 |

We continued the splitting procedure until no further statistically significant divisions could be made, ultimately resulting in 113 classes. Although the underlying ground truth comprises only 16 classes, the 3D classification algorithm failed to distinguish them accurately from the outset. For example, actual class 0 became dispersed across multiple second-iteration classes (C11, C12, C13, C14, C21, and C22, as shown in Table 1). Consequently, additional splits were required to resolve the mixtures introduced by these initial misclassifications. The resulting 113 classes had an average purity of 0.93 (as defined in the legend of Table 1), with class sizes ranging from 41,486 to 145 particles. This demonstrates that our splitting procedure does not merely fragment homogeneous groups into smaller subsets; instead, it applies a statistical criterion for subdivision, independent of class size. For comparison, a single CryoSparc classification into 16 classes yielded a much lower average purity of 0.50.

Fig. 2 displays the *StructMap* projection (Sorzano et al., 2016) of the resulting maps onto a two-dimensional plane. As expected, the projection reveals distinct clusters without smooth transitions, consistent with discrete heterogeneity. Apparent overlaps between some classes are projection artifacts resulting from reducing the original 113*×*113 distance matrix to two dimensions.

**Figure 2:**
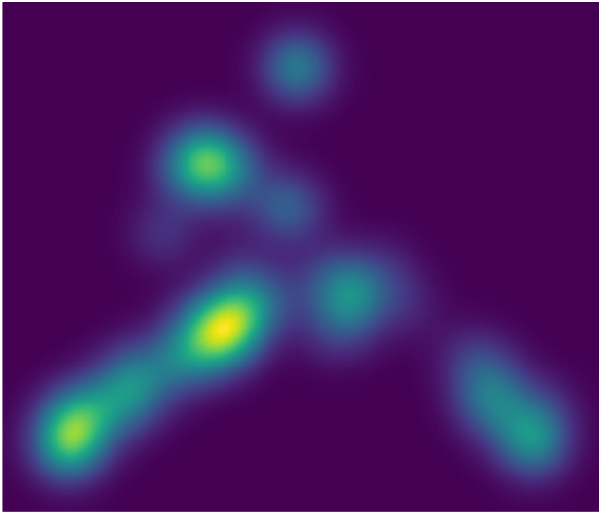
StructMap projection of the Ribosembly dataset. The StructMap provides a low-dimensional representation in which each reconstructed map is positioned according to its similarity to all other maps. In this latent space, more similar maps are placed closer together, while dissimilar maps appear farther apart, providing an intuitive visualization of the structural relationships among classes.

### 3.3 bL17-limited *E. coli* 50S ribosome

The EMPIAR 10841 entry has traditionally been used as a benchmark for discrete 3D classification (Rabuck-Gibbons et al., 2022). The dataset comprises cryo-electron microscopy (cryo-EM) raw movies and processed particle stacks of bL17-limited *Escherichia coli* 50S ribosomal subunit assembly intermediates. This dataset was generated to investigate the structural heterogeneity inherent in ribosome assembly, particularly under conditions where the ribosomal protein bL17 is depleted. This example corresponds to an experimental equivalent of the ribosome assembly simulated dataset.

We initiated our 3D classification using the aligned particles from the dataset. Each iteration involved three independent CryoSparc classifications, identification of statistically distinguishable classes, Fourier reconstruction of each class using Relion’s gridding method, and local angular refinement with Xmipp highres. The original study reported the presence of 41 classes, but since the assignments of individual particles to these classes were not made available in EMPIAR, a direct comparison with our results is not possible. Using our approach, we identified 106 statistically distinguishable classes (see Fig. 3), with particle counts ranging from 400 (see Fig. 4) to 3,400 and an average of approximately 1,300 particles per class. These classes capture both compositional and conformational variability present in the dataset.

**Figure 3:**
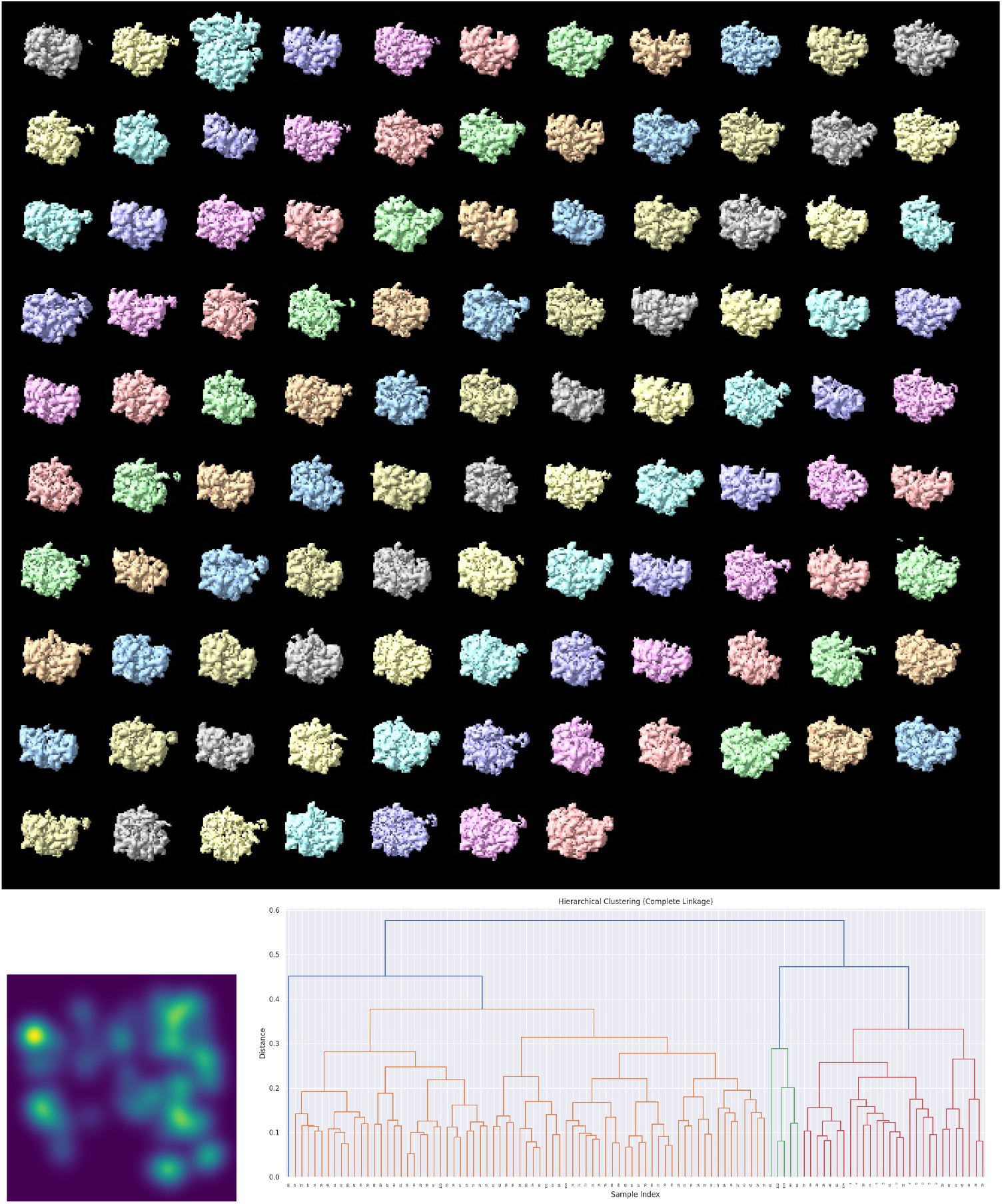
Top: 3D Reconstructions of the particles assigned to each of the classes. Bottom: StructMap projection of the maps and dendrogram representation of their hierarchical clustering. In the dendrogram, maps that are more similar are joined earlier (closer to the leaves), while those that are more dissimilar merge later (closer to the root). The color code is used solely to aid visualization, with groups defined by a default threshold set at 0.7 × max(distance).

**Figure 4:**
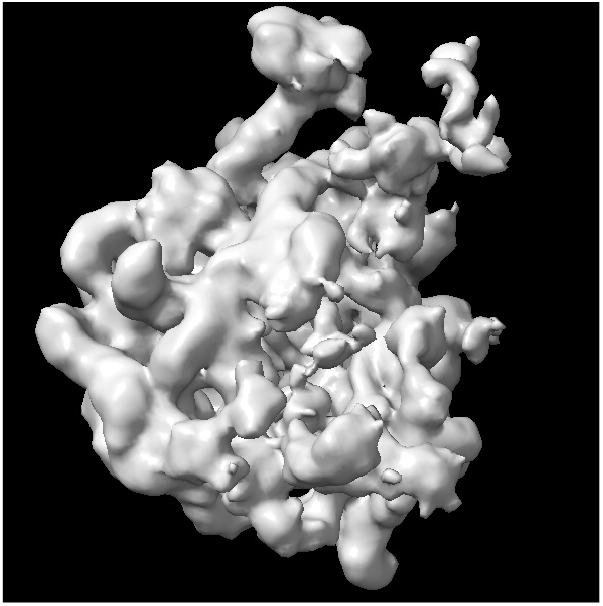
Example reconstruction of the smallest class of the ribosome dataset.

### 3.4 AftD from mycobacteria, Mutant R1389S

The EMPIAR-10391 dataset contains particle images and angular assignments for two conformational states of arabinofuranosyltransferase from a mutant Mycobacterium strain (Tan et al., 2020). According to the original study, the first state comprises 37,814 particles, and the second state includes 68,231. We attempted unsupervised 3D classification into two classes using both *Relion* and *CryoSparc*, each repeated three times. Without masking, neither software could reproducibly separate the two conformations. To improve separation, we applied a spherical mask focused on the region of greatest difference between the two corresponding EMDB maps (IDs: 21600 and 21601).

Using *CryoSparc* for focused classification within the masked region, we repeated the analysis three times and obtained three statistically distinct outcomes: (1) 55,631 particles split 1.5%/98.5%, (2) 46,822 particles split 78%/22%, and (3) 3,592 particles split 13.6%/86.4%. These proportions remained consistent across repetitions, suggesting that the original classification likely included approximately 20% of Class 1 images erroneously assigned to Class 2. To better resolve this heterogeneity, we applied our classification algorithm without angular refinement—justified by the small spatial region distinguishing the classes—and identified 49 statistically distinct classes, with particle counts ranging from 5,369 to 518.

As illustrated in Fig. 5, structural variability within this region is substantial. The *StructMap* projection displays all classes in a two-dimensional space, revealing both densely populated clusters and clearly separated regions. These results demonstrate the ability of standard discrete 3D classification to distinguish structurally distinct particle populations.

**Figure 5:**
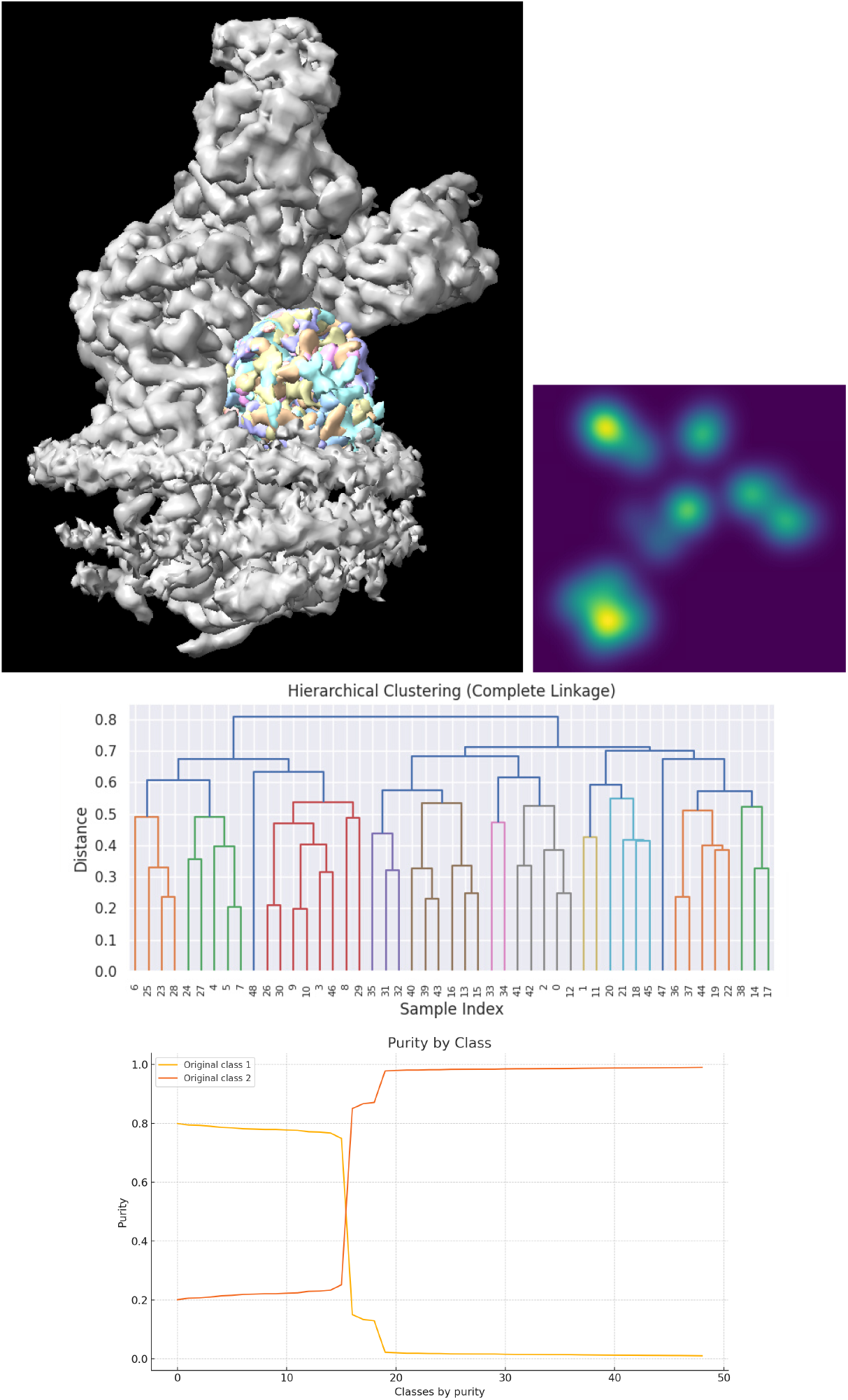
Left: ChimeraX visualization of the 49 classes detected in the EM-PIAR 10391 dataset. Right: StructMap representation of the 49 classes in a two-dimensional projection (the brightness corresponds to multiple classes projecting onto the same coordinate). Middle: Dendrogram representation of the hierarchical clustering of the maps using complete linkage. Bottom: Purity of each of the 49 calculated classes in terms of the original classification deposited in EMPIAR.

While the original two-class scheme captured some of this heterogeneity (as shown in the dendrogram in Fig. 5), it oversimplified the landscape. Many of our classes align closely with the original Class 2, while others correspond mainly to Class 1—though some include particles initially misclassified as Class 2. Apart from this 20% contamination, the original classification underused the potential of discrete classification to uncover the full structural diversity present in the dataset.

## 4 Discussion

Determining the number of meaningful classes in single-particle cryo-EM datasets remains a fundamental and unresolved challenge, particularly in the presence of compositional or conformational heterogeneity. Existing approaches often rely on heuristic criteria—such as class size, map resolution, or visual assessment of class similarity—to guide the classification process. However, such thresholds are inherently arbitrary, lacking a principled statistical foundation, and may lead to inconsistent or non-reproducible results across datasets and users.

In this work, we introduce a novel strategy that incorporates a statistically rigorous criterion—p-values derived from hypothesis testing—to determine whether a given classification step reflects a meaningful structural distinction or merely arises from random partitioning. This framework provides a technically sound basis for deciding when to accept or reject a proposed subdivision, independent of particle count, estimated resolution, or user-defined parameters. By grounding the classification decision in statistical significance, our approach helps mitigate algorithmic bias (when multiple algorithms are used for classification) and subjectivity in the interpretation of classification results. It is essential to emphasize that the assumption of randomness applies only to the null hypothesis, which we aim to disprove. We do not assume that the classification is random—in fact, we think the opposite. By placing the random classification scenario in the null hypothesis, we can use p-values to test whether the observed groupings are statistically distinguishable from randomness, thereby validating their significance.

A critical advantage of our method is that it does not inherently favor the formation of small classes. On the contrary, we have demonstrated its flexibility across a broad spectrum of class sizes: in some datasets, statistically distinguishable classes contained several tens of thousands of particles, while in others, valid classes were composed of just a few hundred. This adaptability underscores the data-driven nature of our approach, which bases subdivisions on reproducibility rather than on fixed size or resolution thresholds.

Our results also highlight the intrinsic instability of current discrete 3D classification algorithms. Even when using the same input data and classification parameters, repeated runs can yield divergent class assignments—a consequence of noise, alignment uncertainties, and algorithmic non-determinism. Rather than avoiding this variability, our method embraces it by explicitly testing the reproducibility of classifications. Subsets of particles that consistently appear together across independent runs are considered statistically significant, while unstable groupings are rejected. In this way, the instability of 3D classification becomes a measurable property that can be harnessed to improve confidence in the final set of classes.

Despite its advantages, our algorithm is ultimately constrained by the capabilities of the underlying 3D classification algorithm used to detect structural differences. The statistical framework we propose relies on the reproducibility of these classifications; however, if the 3D classifier fails to resolve an actual structural distinction, due, for example, to low signal-to-noise ratio (SNR) or subtle conformational differences, then no statistical test can recover that information. In this sense, the sensitivity of our method is fundamentally limited by the detectability of differences in the data itself. When structural variations fall below the resolution threshold imposed by noise or insufficient particle count, even repeated classification and hypothesis testing may fail to identify meaningful subdivisions. Thus, while our algorithm adds robustness to the interpretation of 3D classification results, it cannot compensate for the inherent limitations of the data or the algorithms used to process it.

In conclusion, our approach provides a principled framework for identifying the number of structurally meaningful classes in cryo-EM datasets. By replacing arbitrary thresholds with statistically validated decisions, it enhances the interpretability and reproducibility of 3D classification, offering a promising direction for future developments in cryo-EM heterogeneity analysis.

## Acknowledgements

The authors acknowledge financial support from the Ministry of Science, Innovation and Universities (BDNS No. 716450) to Instruct-Spain, as part of Spain’s participation in Instruct–ERIC, the European Strategic Infrastructure Project (ESFRI) in Structural Biology. This work was also supported by Project PID2022-136594NB-I00, funded by MICIU/AEI/10.13039/501100011033 and the European Regional Development Fund (ERDF) under the slogan “A way of making Europe”, as well as by the Spanish State Research Agency (AEI/10.13039/501100011033) through the “Severo Ochoa” Centres of Excellence in R&D Programme [Project CEX2023-001386-S]. The Comunidad Autónoma de Madrid provided additional support through Project S2022/BMD-7232, and by the European Union under the Horizon Europe Programme through the following projects: Fragment-Screen [Project 101094131], Harmony EuroHPC [Project 101163317], EOSC Beyond [Project 101131875], and EnLaCES [Project 101024130]. The project that gave rise to these results received the support of a fellowship from “la Caixa” Foundation (ID 100010434). The fellowship code is LCF/BQ/DR24/1208003.

## References

Abrishami, V., Bilbao-Castro, J. R., Vargas, J., Marabini, R., Carazo, J. M., and Sorzano, C. O. S. (2015). A fast iterative convolution weighting approach for gridding-based direct fourier three-dimensional reconstruction with correction for the contrast transfer function. Ultramicroscopy, 157:79–87.

Aiyer, S., Baldwin, P. R., Tan, S. M., Shan, Z., Oh, J., Mehrani, A., Bowman, M. E., Louie, G., Passos, D. O., Djorjevic-Marquardt, S., Mietzsch, M., Hull, J. A., Hoshika, S., Barad, B. A., Grotjahn, D. A., McKenna, R., Agbandje-McKenna, M., Benner, S. A., Noel, J. A. P., Wang, D., Tan, Y. Z., and Lyumkis, D. (2024). Overcoming resolution attenuation during tilted cryo-EM data collection. Nature communications, 15(1):389.

Gomez-Blanco, J., Kaur, S., Strauss, M., and Vargas, J. (2022). Hierarchical autoclassification of cryo-EM samples and macromolecular energy landscape determination. Computer Methods and Programs in Biomedicine, 216:106673.

Jeon, M., Raghu, R., Astore, M., Woollard, G., Feathers, J., Kaz, A., Hanson, S., Cossio, P., and Zhong, E. (2024). CryoBench: Diverse and challenging datasets for the heterogeneity problem in cryo-EM. Advances in Neural Information Processing Systems, 37:89468–89512.

Kimanius, D. and Schwab, J. (2024). Confronting heterogeneity in cryogenic electron microscopy data: Innovative strategies and future perspectives with data-driven methods. Current Opinion in Structural Biology, 86:102815.

Rabuck-Gibbons, J. N., Lyumkis, D., and Williamson, J. R. (2022). Quantitative mining of compositional heterogeneity in cryo-EM datasets of ribosome assembly intermediates. Structure, 30:498–509.e4.

Sorzano, C. O. S., Alvarez-Cabrera, A. L., Kazemi, M., Carazo, J. M., and Jonić, S. (2016). Structmap: Elastic distance analysis of electron microscopy maps for studying conformational changes. Biophys J, 110:1753–1765.

Sorzano, C. O. S., Jiménez, A., Mota, J., Vilas, J. L., Maluenda, D., Martínez, M., Ramírez-Aportela, E., Majtner, T., Segura, J., Sánchez-García, R., Rancel, Y., Del Caño, L., Conesa, P., Melero, R., Jonic, S., Vargas, J., Cazals, F., Freyberg, Z., Krieger, J., Bahar, I., Marabini, R., and Carazo, J. M. (2019). Survey of the analysis of continuous conformational variability of biological macromolecules by electron microscopy. Acta crystallographica. Section F, Structural biology communications, 75:19–32.

Sorzano, C. O. S., Jiménez-Moreno, A., Maluenda, D., Martinez, M., Ramirez-Aportela, E., Melero, R., Cuervo, A., Conesa, J., Filipovic, J., Conesa, P., del Caño, L., Fonseca, Y. C., Jiménez-de la Morena, J., Losana, P., Sánchez-García, R., Strelak, D., Fernández-Giménez, E., de Isidro-Gómez, F., Her-reros, D., Vilas, J. L., Marabini, R., and Carazo, J. M. (2022). On bias, variance, overfitting, gold standard and consensus in single particle analysis by cryo-electron microscopy. Acta Crystallographica Section D, D78:410–423.

Sorzano, C. O. S., Vargas, J., de la Rosa-Trevin, J.M., Jimenez, A., Maluenda, D., Melero, R., Martinez, M., Ramirez-Aportela, E., Conesa, P., Vilas, J. L., Marabini, R., and Carazo, J. M. (2018). A new algorithm for high-resolution reconstruction of single particles by electron microscopy. J. Structural Biology, 204:329–337.

Tan, Y. Z., Zhang, L., Rodrigues, J., Zheng, R. B., Giacometti, S. I., Rosário, A. L., Kloss, B., Dandey, V. P., Wei, H., Brunton, R., et al. (2020). Cryo-EM structures and regulation of arabinofuranosyltransferase aftd from mycobacteria. Molecular cell, 78(4):683–699.

Tang, W. S., Zhong, E. D., Hanson, S. M., Thiede, E. H., and Cossio, P. (2023). Conformational heterogeneity and probability distributions from single-particle cryo-electron microscopy. Current Opinion in Structural Biology, 81:102626.

Tibshirani, R., Walther, G., and Hastie, T. (2001). Estimating the number of clusters in a data set via the gap statistic. J. royal statistical society: series b (statistical methodology), 63(2):411–423.

Toader, B., Sigworth, F. J., and Lederman, R. R. (2023). Methods for cryo-EM single particle reconstruction of macromolecules having continuous heterogeneity. J. Molecular Biology, 435(9):168020.

Zhou, Y., Moscovich, A., and Bartesaghi, A. (2022). Data-driven determination of number of discrete conformations in single-particle cryo-EM. Computer Methods and Programs in Biomedicine, 221:106892.

